# Time restricted feeding with or without ketosis influences metabolism-related gene expression in a tissue-specific manner in aged rats

**DOI:** 10.1101/2024.12.19.629431

**Authors:** Sarah Ding, Anisha Banerjee, Sara N. Burke, Abbi R. Hernandez

## Abstract

Many of the ‘hallmarks of aging’ involve alterations in cellular and organismal metabolism. One pathway with the potential to impact several traditional markers of impaired function with aging is the PI3K/AKT metabolic pathway. Regulation of this pathway includes many aspects of cellular function, including protein synthesis, proliferation and survival, as well as many downstream targets, including mTOR and FOXOs. Importantly, this pathway is pivotal to the function of every organ system in the human body. Thus, we investigated the expression of several genes along this pathway in multiple organs, including the brain, liver and skeletal muscle, in aged subjects that had been on different experimental diets to regulate metabolic function since mid-life. Specifically, rats were fed a control ad lib diet (AL), a time restricted feeding diet (cTRF), or a time restricted feeding diet with ketogenic macronutrients (kTRF) for the majority of their adult lives (from 8-25 months). We previously reported that regardless of macronutrient ratio, TRF-fed rats in both macronutrient groups required significantly less training to acquire a biconditional association task than their ad lib fed counterparts. The current experiments expand on this work by quantifying metabolism-related gene expression across tissues and interrogating for potential relationships with cognitive performance. AKT expression was significantly reduced in kTRF fed rats within liver and muscle tissue. However, AKT expression within the perirhinal cortex (PER) was higher in kTRF rats with the best cognitive performance. Within CA3, higher levels of FOXO1 gene expression correlated with poorer cognitive performance in ad libitum fed rats. Together, these data demonstrate diet- and tissue-specific alterations in metabolism-related gene expression and their correlation with cognitive status.

## Introduction

Aging is associated with changes in metabolic function at every level, from cellular [1] to organismal [2]. Unsurprisingly, this extends to one of the most pivotal pathways in cellular metabolism - the PI3K/AKT pathway. This pathway is involved in many aspects of cellular function, including protein synthesis, proliferation and survival. The activation of AKT results in possible phosphorylation of a wide range of targets, many of which are critical for metabolic function, including the mechanistic target of rapamycin (mTOR), glycogen synthase kinase 3 (GSK3) and forkhead box O (FOXO). Interestingly, reducing activation of this pathway can extend the lifespan of species at several organismal levels [3], though full ablation of this pathway results in deleterious effects such as cardiomyopathy [4] and hepatocellular carcinoma [5].

Aberrant activity within this pathway is associated with many age-associated disease states, including diabetes, cancer and neurodegeneration [6]. In fact, heightened activity within this pathway is an early feature of Alzheimer’s disease (AD) [7], likely due to increased activation by insulin like growth factor (IGF-1R) and insulin receptors (IR) [8]. Moreover, changes in signaling along this pathway can profoundly impact cognitive function, due to its key role in long term potentiation and depression (LTP/D) [9].

Proper regulation of the PI3K/AKT pathway is not only necessary for proper brain function but plays a key role throughout the entire body. Within muscle tissue, this pathway plays many roles, including the prevention of muscle atrophy and the maintenance of glucose homeostasis [10], due to its pivotal role in the mediation of insulin-stimulated glucose uptake [11]. Within the liver, this pathway is necessary for the regulation of glucose metabolism, especially FOXO1 [12]. Overexpression of AKT in liver tissue can lead to hypoglycemia and lipid accumulation [12].

One method with the potential to significantly alter expression of this pathway across tissues is through dietary intervention. Two methods in particular have been proposed to benefit metabolic health and potentially regulate this pathway: ketogenic diets (KDs) and time restricted feeding (TRF; also known as intermittent fasting and time restricted eating). KDs are high in fat and sufficiently restricted in carbohydrate content such that there is a switch from glucose to ketone bodies (produced from fat) as the primary fuel source for the tricarboxylic acid cycle. While historically these diets are primarily used for the treatment of refractory epilepsy [13] and glucose transporter 1 deficiency [14], they have gained popularity for the treatment of many age-related disease states including AD [15] and cancer [16]. While aged subjects may be delayed in this metabolic switch [17], they are capable of nutritional ketosis and benefitting from positive changes to their overall metabolic state [18]. KDs are also typically associated with loss of body fat and improved glycemic control [19]. Unlike KDs, TRFs do not require a shift in macronutrient composition of the diet, but may also be associated with a glycolysis to ketosis switch if the fasting window is sufficient [20]. Instead, the time window during which calories are consumed is restricted. This type of feeding paradigm is also linked with improved body composition [21] and glucose homeostasis [22]. We therefore wanted to explore the effects of long-term TRF with and without ketogenic macronutrient composition on PI3K/AKT pathway related gene expression across multiple tissue types (brain, liver and muscle) in a cohort of aged subjects. Because data regarding the safety and efficacy of these dietary paradigms long term in humans is not feasible to acquire, we investigated the effects of ad lib feeding, control with TRF (cTRF) and KD with TRF (kTRF) in aged rats consuming these diets for over half their lives. Gene expression was specific to both dietary intervention and tissue type. Moreover, even within the brain, the effects varied between regions. Our previous work demonstrated that both KDs [23] and TRF [24] can improve age-related impairments on associative task performance [25], which depends on proper functioning across the prefrontal (PFC) and perirhinal (PER) cortices [26]. Herein, there were significant correlations with gene expression and cognitive performance on this associative learning task.

## Methods

### Subjects and interventions

33 male Fisher 344 x Brown Norway rats were housed individually and maintained on a reverse 12-hr light/12-hr dark cycles so that all feeding and cognitive testing occurred during the dark/active phase of the diurnal cycle. Cognitive function and details regarding behavioral training and testing from the rats utilized herein have been previously reported [24]. Rats were divided into three diet groups: fed 51 kCal of a standard control diet once daily in a time restricted feeding (TRF) paradigm (cTRF;n=10), fed 51 kCal of a ketogenic TRF diet (kTRF; n=10), and a group fed standard rat chow *ad libitum* (AL; n=13). All rats were fed these diets from 8-21 months, at which point their food intake was modestly restricted to encourage participation in the behavioral tasks previously described.^18^ The ketogenic diet was a high fat/low carbohydrate diet with MCT oil (76% fat, 4% carbohydrates, and 20% protein), and the control diet was 16% fat, 65% carbohydrates, and 19% protein. The ketogenic and control TRF rats were calorie and micronutrient matched. It is also important to note that the rats were not calorically restricted, as they continued to show increases in weight throughout their lives. For complete details of this study, please refer to the original publication.^18^

After completing behavioral testing (at 24-26 months of age), liver, muscle (tibialis anterior) and brain tissue were collected. Five regions within the temporal lobe were collected: the perirhinal cortex (PER), entorhinal cortex (ENT), as well as the dentate gyrus (DG), CA1 and CA3 of the hippocampus. Tissues were flash frozen on dry ice and preserved at -80C°.

### Reverse Transcription and PCR Expression Assay

RNA was isolated with a cold TRIzol (Thermo Fisher Scientific) extraction, or for samples <30mg, via PureLink RNA Micro Kit (Invitrogen). Total RNA was quantified with a NanoDrop UV Visible Spectrophotometer machine. cDNA was synthesized using a Super Script VILO cDNA synthesis kit (Thermo Fisher Scientific) according to the instructions provided. Samples were diluted and stored at -20C° until use. Relative gene expression of each tissue was determined using a QuantStudio 5 System PCR machine with the target gene primers listed in Table 1, diluted to 1X. Actin beta was used as a housekeeping gene for each target gene. The fast PCR amplification reaction procedure consisted of the following steps: an initial denaturation step that lasted for 20 seconds at 95° C, followed by 40 cycles of denaturation at 95° C for 1 second and extension at 60° C for 20 seconds. Ct values, or cycle threshold values, were recorded for each gene and the 2^-ΔΔCt^ method was used to calculate the relative fold gene expression of samples.

**Table 1.**
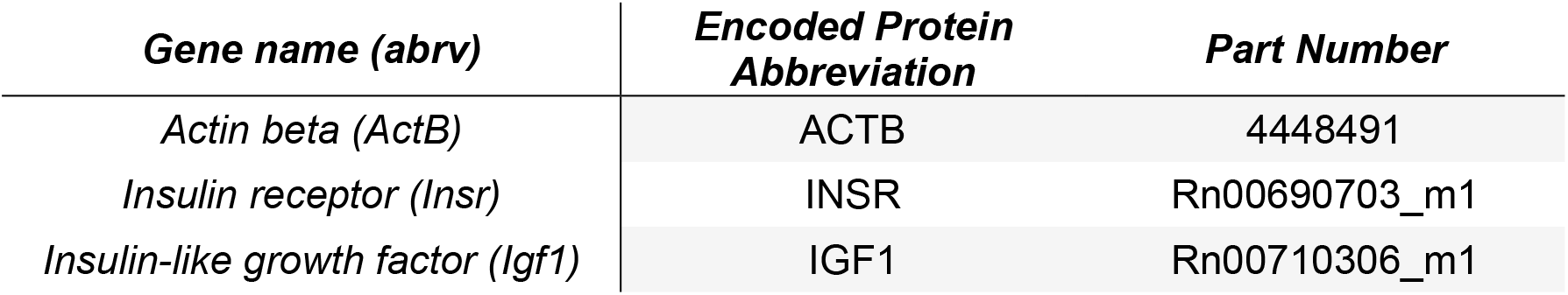
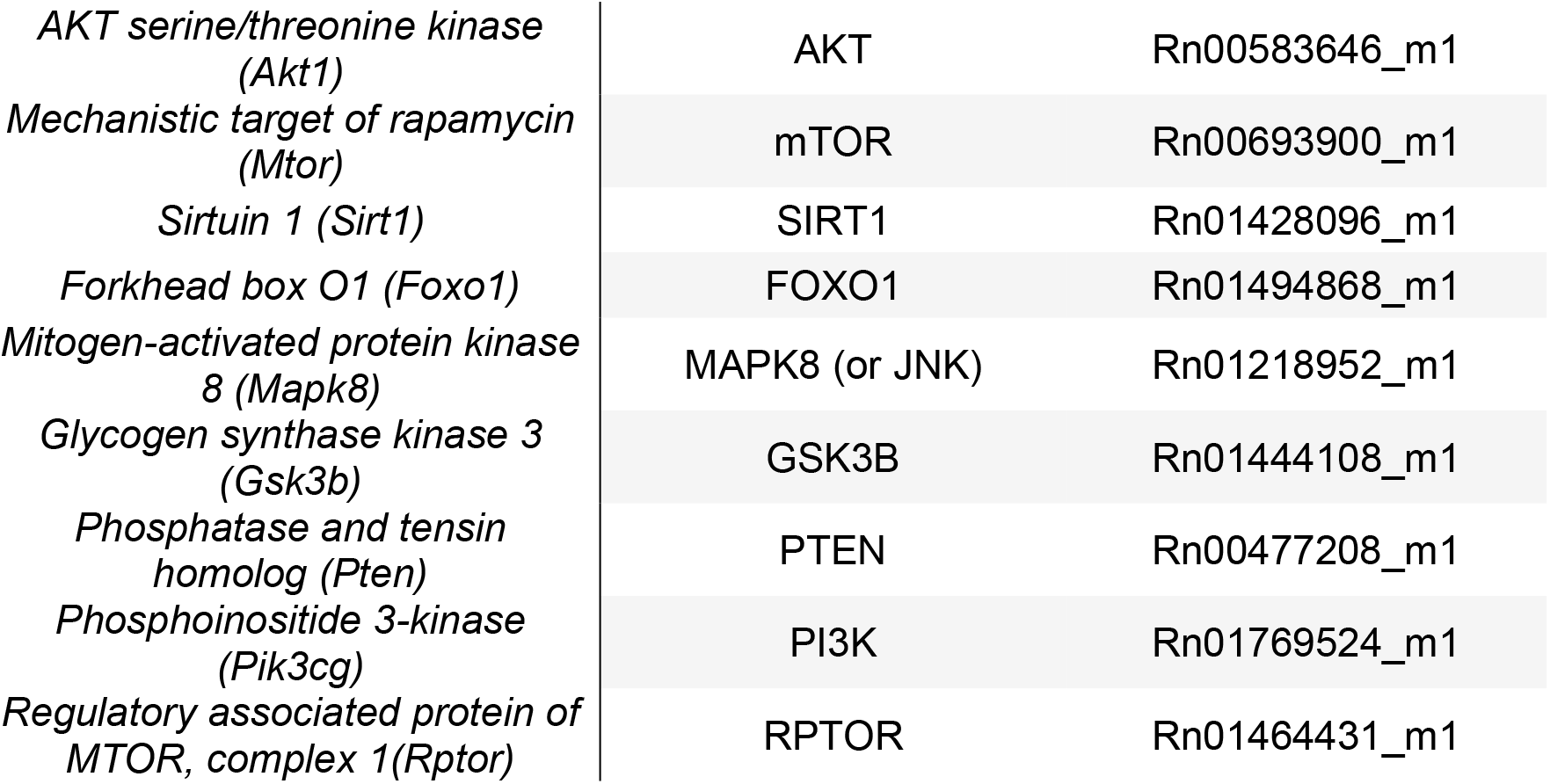
PCR Primers, all sourced from Thermo Fisher Scientific.

### Statistical Analysis

All quantitative data are expressed as group means ± the standard error of the mean (SEM) relative to the expression of the same gene within the same tissue from rats in the control group. Data were first analyzed with a Linear Mixed Model (LMM) to increase accuracy of reporting by accounting for the potential correlation induced by the repeated measure of gene expression across different tissue types. For this model, each rat was considered a repeated random effect, and diet group and tissue type were considered fixed effects. Extreme outliers, identified as any data greater than 3 times above or below the interquartile range, were excluded. A type 3 test was then used to investigate main effects of tissue type and diet group for each gene, as well as the interaction between these two factors. Post-hoc analyses were conducted via pair-wise comparisons. Due to the large number of comparisons presented herein, pairwise comparisons corrected within each tissue using Tukey tests are presented in the results section. However, for complete transparency of data we also present an unadjusted p-value, p-value adjusted with Tukey’s test within each tissue type and p-value adjusted using Tukey within each tissue as well as false discovery rate (FDR) between tissues (double adjusted) in table 2. Correlation analyses were performed using the Hmisc^26^ and corrplot R packages^27^ for each feeding paradigm (TRF or AL) with expression of each gene and number of incorrect trials until reaching criterion on an object-place paired association task performed in old age. The null hypothesis was rejected at the level of *p* > 0.05, unless otherwise stated.

**Table 2.**
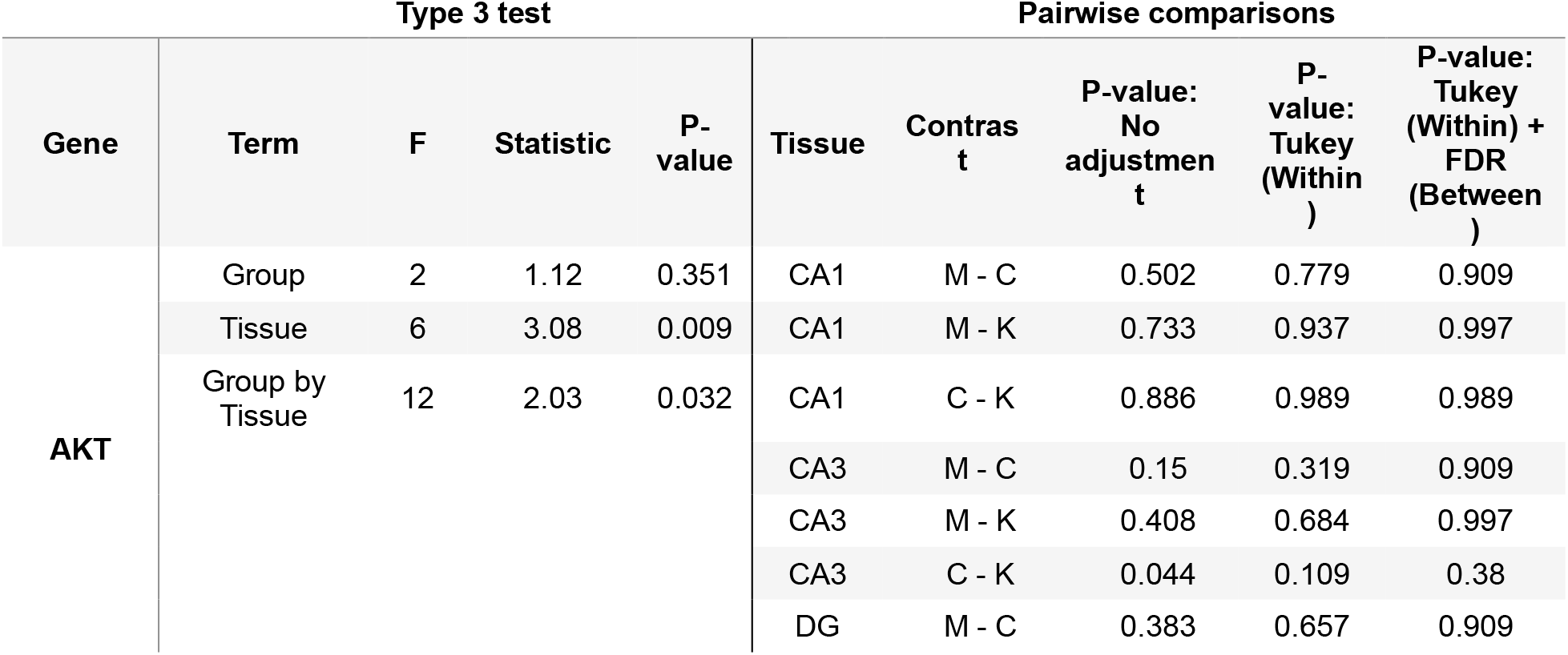
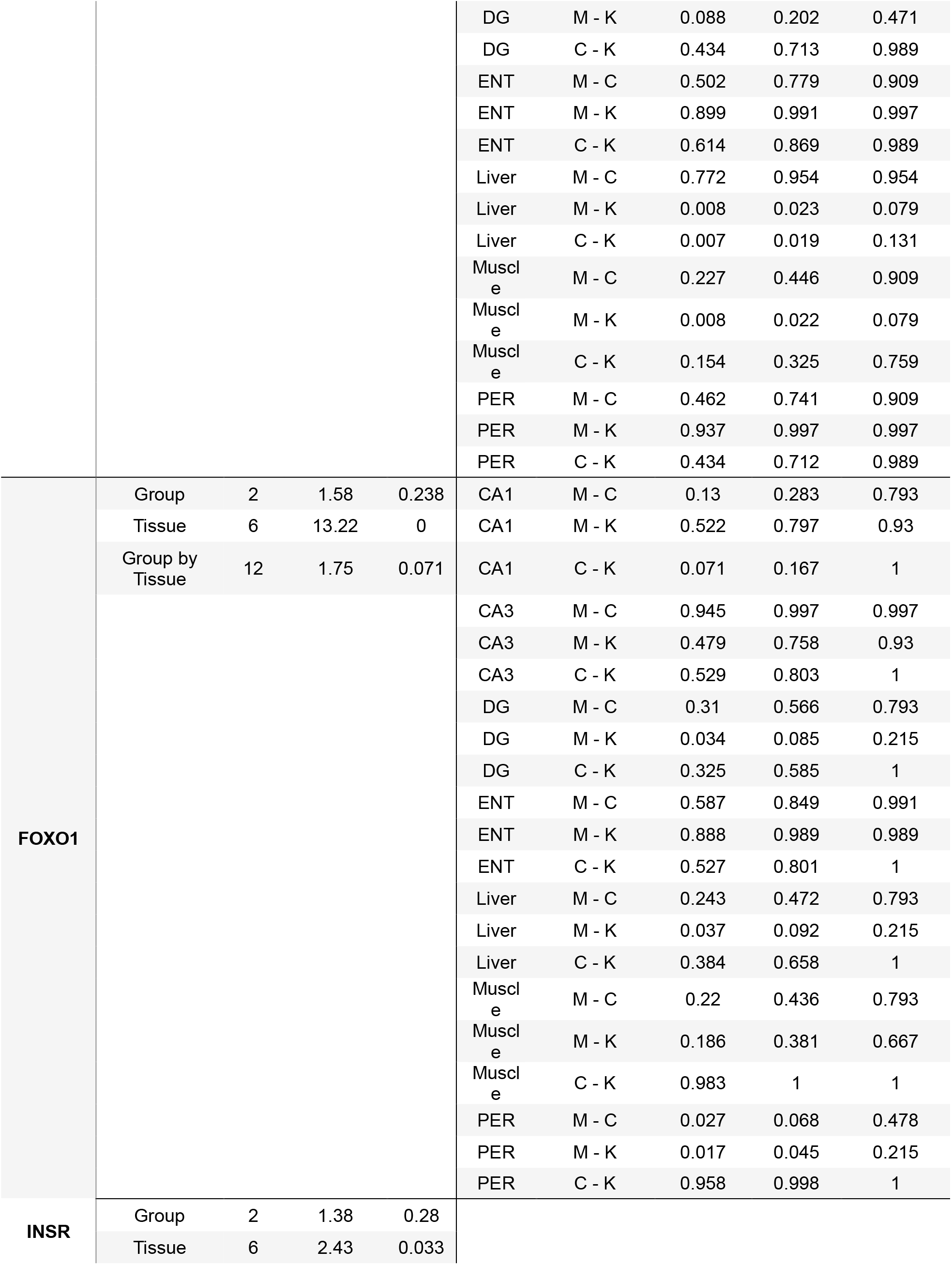
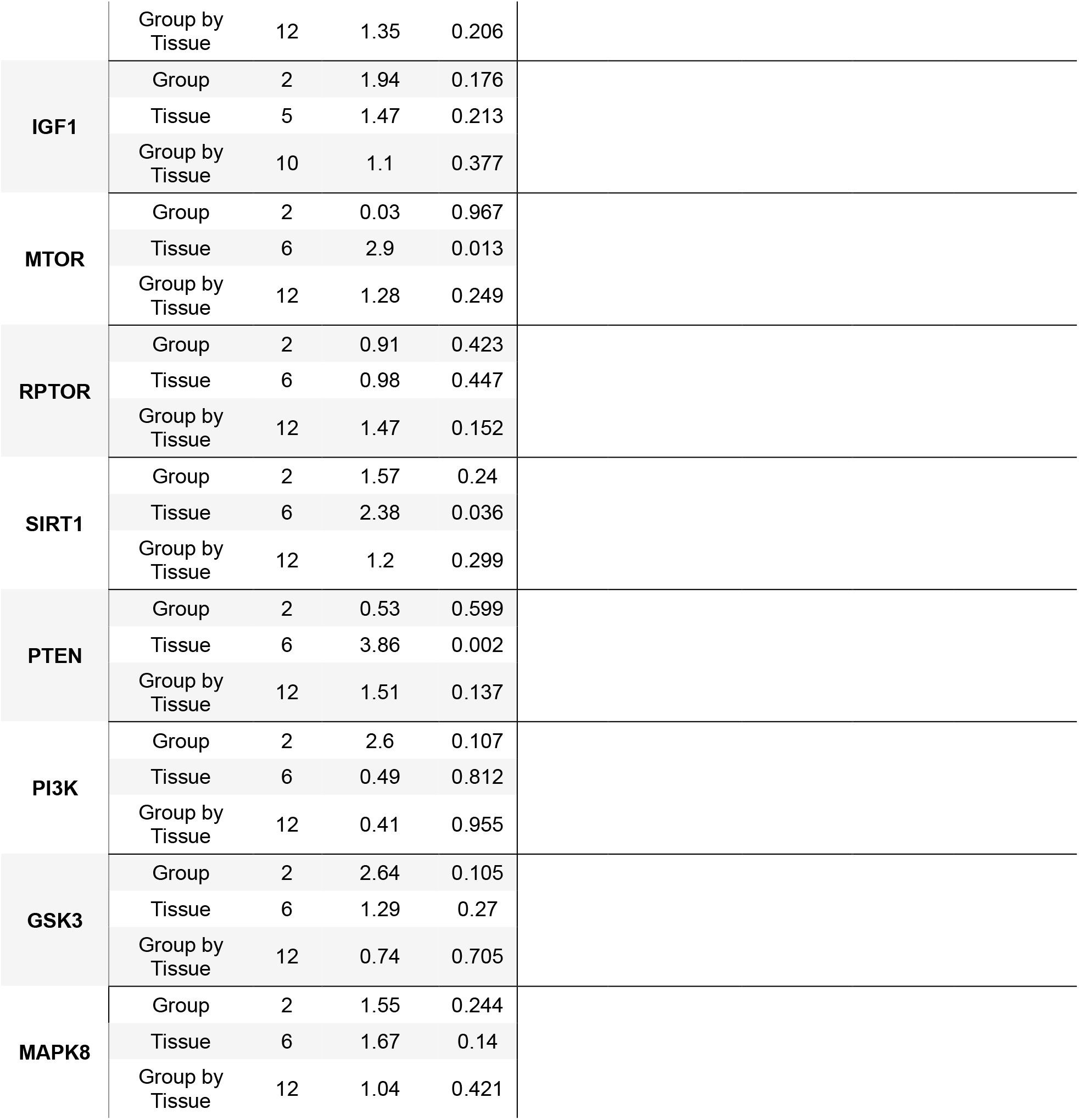
Statistical evaluation of gene expression across diets and tissues with unadjusted p-values, p-values corrected with Tukey within subjects and p-values double corrected with both Tukey within subjects and false discovery rate (FDR) between subjects.

| Type 3 test |  |  |  |  | Pairwise comparisons |  |  |  |  |
| --- | --- | --- | --- | --- | --- | --- | --- | --- | --- |
| Gene | Term | F | Statistic | P-value | Tissue | Contrast | P-value: No adjustment | P-value: Tukey (Within) | P-value: Tukey (Within) + FDR (Between) |
| AKT | Group | 2 | 1.12 | 0.351 | CA1 | M - C | 0.502 | 0.779 | 0.909 |
|  | Tissue | 6 | 3.08 | 0.009 | CA1 | M - K | 0.733 | 0.937 | 0.997 |
|  | Group by Tissue | 12 | 2.03 | 0.032 | CA1 | C - K | 0.886 | 0.989 | 0.989 |
|  |  |  |  |  | CA3 | M - C | 0.15 | 0.319 | 0.909 |
|  |  |  |  |  | CA3 | M - K | 0.408 | 0.684 | 0.997 |
|  |  |  |  |  | CA3 | C - K | 0.044 | 0.109 | 0.38 |
|  |  |  |  |  | DG | M - C | 0.383 | 0.657 | 0.909 |
|  |  |  |  |  | DG | M - K | 0.088 | 0.202 | 0.471 |
|  |  |  |  |  | DG | C - K | 0.434 | 0.713 | 0.989 |
|  |  |  |  |  | ENT | M - C | 0.502 | 0.779 | 0.909 |
|  |  |  |  |  | ENT | M - K | 0.899 | 0.991 | 0.997 |
|  |  |  |  |  | ENT | C - K | 0.614 | 0.869 | 0.989 |
|  |  |  |  |  | Liver | M - C | 0.772 | 0.954 | 0.954 |
|  |  |  |  |  | Liver | M - K | 0.008 | 0.023 | 0.079 |
|  |  |  |  |  | Liver | C - K | 0.007 | 0.019 | 0.131 |
|  |  |  |  |  | Muscle | M - C | 0.227 | 0.446 | 0.909 |
|  |  |  |  |  | Muscle | M - K | 0.008 | 0.022 | 0.079 |
|  |  |  |  |  | Muscle | C - K | 0.154 | 0.325 | 0.759 |
|  |  |  |  |  | PER | M - C | 0.462 | 0.741 | 0.909 |
|  |  |  |  |  | PER | M - K | 0.937 | 0.997 | 0.997 |
|  |  |  |  |  | PER | C - K | 0.434 | 0.712 | 0.989 |
| FOXO1 | Group | 2 | 1.58 | 0.238 | CA1 | M - C | 0.13 | 0.283 | 0.793 |
|  | Tissue | 6 | 13.22 | 0 | CA1 | M - K | 0.522 | 0.797 | 0.93 |
|  | Group by Tissue | 12 | 1.75 | 0.071 | CA1 | C - K | 0.071 | 0.167 | 1 |
|  |  |  |  |  | CA3 | M - C | 0.945 | 0.997 | 0.997 |
|  |  |  |  |  | CA3 | M - K | 0.479 | 0.758 | 0.93 |
|  |  |  |  |  | CA3 | C - K | 0.529 | 0.803 | 1 |
|  |  |  |  |  | DG | M - C | 0.31 | 0.566 | 0.793 |
|  |  |  |  |  | DG | M - K | 0.034 | 0.085 | 0.215 |
|  |  |  |  |  | DG | C - K | 0.325 | 0.585 | 1 |
|  |  |  |  |  | ENT | M - C | 0.587 | 0.849 | 0.991 |
|  |  |  |  |  | ENT | M - K | 0.888 | 0.989 | 0.989 |
|  |  |  |  |  | ENT | C - K | 0.527 | 0.801 | 1 |
|  |  |  |  |  | Liver | M - C | 0.243 | 0.472 | 0.793 |
|  |  |  |  |  | Liver | M - K | 0.037 | 0.092 | 0.215 |
|  |  |  |  |  | Liver | C - K | 0.384 | 0.658 | 1 |
|  |  |  |  |  | Muscle | M - C | 0.22 | 0.436 | 0.793 |
|  |  |  |  |  | Muscle | M - K | 0.186 | 0.381 | 0.667 |
|  |  |  |  |  | Muscle | C - K | 0.983 | 1 | 1 |
|  |  |  |  |  | PER | M - C | 0.027 | 0.068 | 0.478 |
|  |  |  |  |  | PER | M - K | 0.017 | 0.045 | 0.215 |
|  |  |  |  |  | PER | C - K | 0.958 | 0.998 | 1 |
| INSR | Group | 2 | 1.38 | 0.28 |  |  |  |  |  |
|  | Tissue | 6 | 2.43 | 0.033 |  |  |  |  |  |

|  |  |  |  |  |
| --- | --- | --- | --- | --- |
|  | Group by Tissue | 12 | 1.35 | 0.206 |
| IGF1 | Group | 2 | 1.94 | 0.176 |
|  | Tissue | 5 | 1.47 | 0.213 |
|  | Group by Tissue | 10 | 1.1 | 0.377 |
| MTOR | Group | 2 | 0.03 | 0.967 |
|  | Tissue | 6 | 2.9 | 0.013 |
|  | Group by Tissue | 12 | 1.28 | 0.249 |
| RPTOR | Group | 2 | 0.91 | 0.423 |
|  | Tissue | 6 | 0.98 | 0.447 |
|  | Group by Tissue | 12 | 1.47 | 0.152 |
| SIRT1 | Group | 2 | 1.57 | 0.24 |
|  | Tissue | 6 | 2.38 | 0.036 |
|  | Group by Tissue | 12 | 1.2 | 0.299 |
| PTEN | Group | 2 | 0.53 | 0.599 |
|  | Tissue | 6 | 3.86 | 0.002 |
|  | Group by Tissue | 12 | 1.51 | 0.137 |
| PI3K | Group | 2 | 2.6 | 0.107 |
|  | Tissue | 6 | 0.49 | 0.812 |
|  | Group by Tissue | 12 | 0.41 | 0.955 |
| GSK3 | Group | 2 | 2.64 | 0.105 |
|  | Tissue | 6 | 1.29 | 0.27 |
|  | Group by Tissue | 12 | 0.74 | 0.705 |
| MAPK8 | Group | 2 | 1.55 | 0.244 |
|  | Tissue | 6 | 1.67 | 0.14 |
|  | Group by Tissue | 12 | 1.04 | 0.421 |

## Results

### The effects of dietary intervention on gene expression are tissue specific

Gene expression data were first analyzed with a Linear Mixed Model (LMM) running a Type 3 Test for each gene. Gene expression differed across tissue type for the following genes: Akt1 (p = 0.009), Insr (p = 0.033), Mtor (p = 0.013), Sirt1 (p = 0.036), Foxo1 (p < 0.001), Pten (p = 0.002). However, the only significant interaction between tissue type and diet group was for Akt1 (p = 0.03), as well as a trend towards an interaction for Foxo1 (p = 0.07).

Pairwise comparisons by diet were used to interrogate the extent that Akt1 and Foxo1 expression within individual tissue types significantly varied by diet condition (**Figure 1**). Within the liver, Akt1 expression was significantly lower following the ketogenic diet than the ad lib control (p = 0.023) and cTRF (p = 0.019) groups. Within the muscle, kTRF rats similarly had lower expression than the ad lib control fed rats (p = 0.022). Pairwise comparisons for Foxo1 expression between diet groups showed that expression was significantly lower in kTRF rats relative to ad lib controls within the PER only (p = 0.045). Similarly, there was a strong trend towards decreased expression in cTRF fed rats relative to ad lib controls as well (p = 0.068), indicating this effect may be due to TRF rather than macronutrient ratio. While no other pairwise interactions were significantly different following adjustment for multiple comparisons, p-values are presented in **table 2**, using all 3 ways described in the methods, for data transparency.

**Figure 1.**
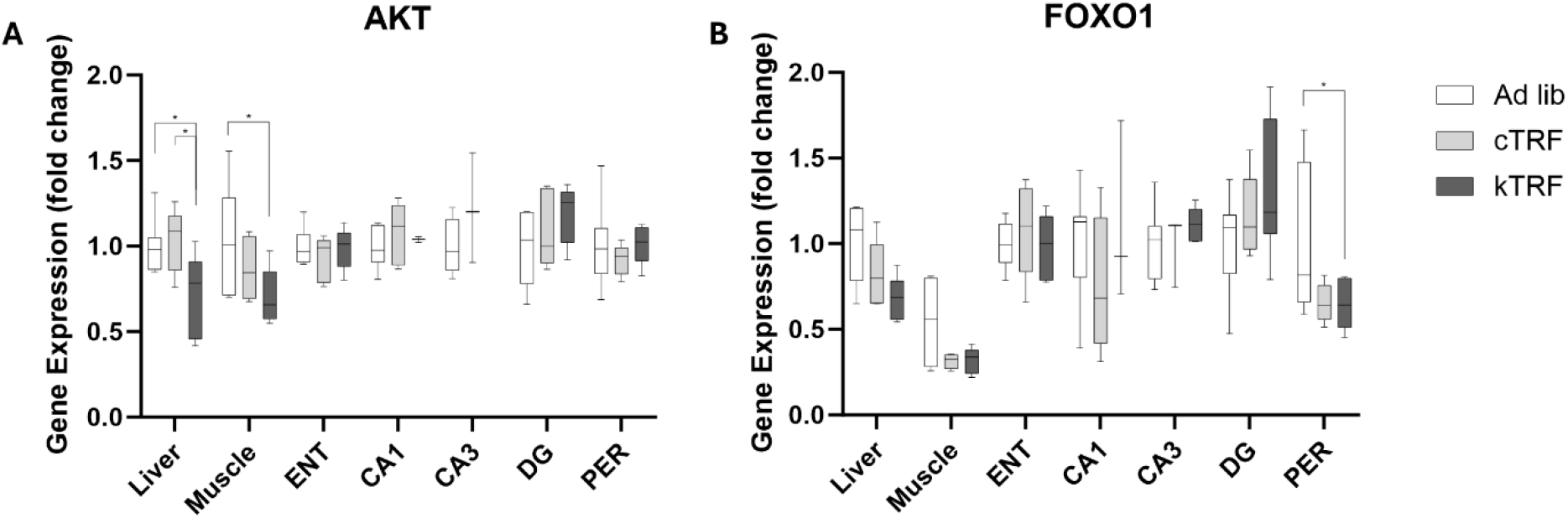
Gene expression is significantly altered by dietary intervention in a tissue-specific manner. (A) Akt1 expression was significantly lowered in kTRF rats relative to both control diet conditions within the liver and relative to ad lib control diet within the muscle. Levels in brain remained unaffected. (B) Conversely, Foxo1 significantly lowered gene expression within the perirhinal cortex (PER) of the brain and not within peripheral tissues.

### Tissue-specific gene expression is associated with behavioral performance in a diet-based manner

Cognitive performance in these rats was assessed with the object place paired associated (OPPA) task and these data have previously been published [24]. Briefly, the OPPA task requires rats to perform a simple pairwise object comparison in a two-arm arm maze (e.g., owl figurine versus turtle figurine) for a food reward. However, the correct choice of objects differs by location on the maze with the owl being the correct choice in one arm and the turtle being the correct choice in the opposite arm. Both TRF fed groups, regardless of macronutrient ratio, learned the object-in-place rule to a criterion performance of above 85% correct in fewer training trials than ad lib control fed rats. OPPA performance was specifically quantified as the number of errors made prior to reaching criterion, with a higher value indicating a worse performance. Therefore, the relationship between the genes significantly altered across diet and tissue and the number of incorrect trials required to perform the object-in-place rule to criterion was interrogated individually for each gene with a significant group by tissue interaction. When analyzed across all rats, regardless of diet group, there was a significant correlation between Akt1 expression within the PER and behavioral performance (r = -0.55; p = 0.02; **Figure 2A**). There was also a trend towards an association between OPPA task performance and liver Akt1 expression (r = 0.46; p = 0.07), but Akt1 expression in all other tissues did not show a correlation with OPPA performance (p > 0.31 for all remaining comparisons).

**Figure 2.**
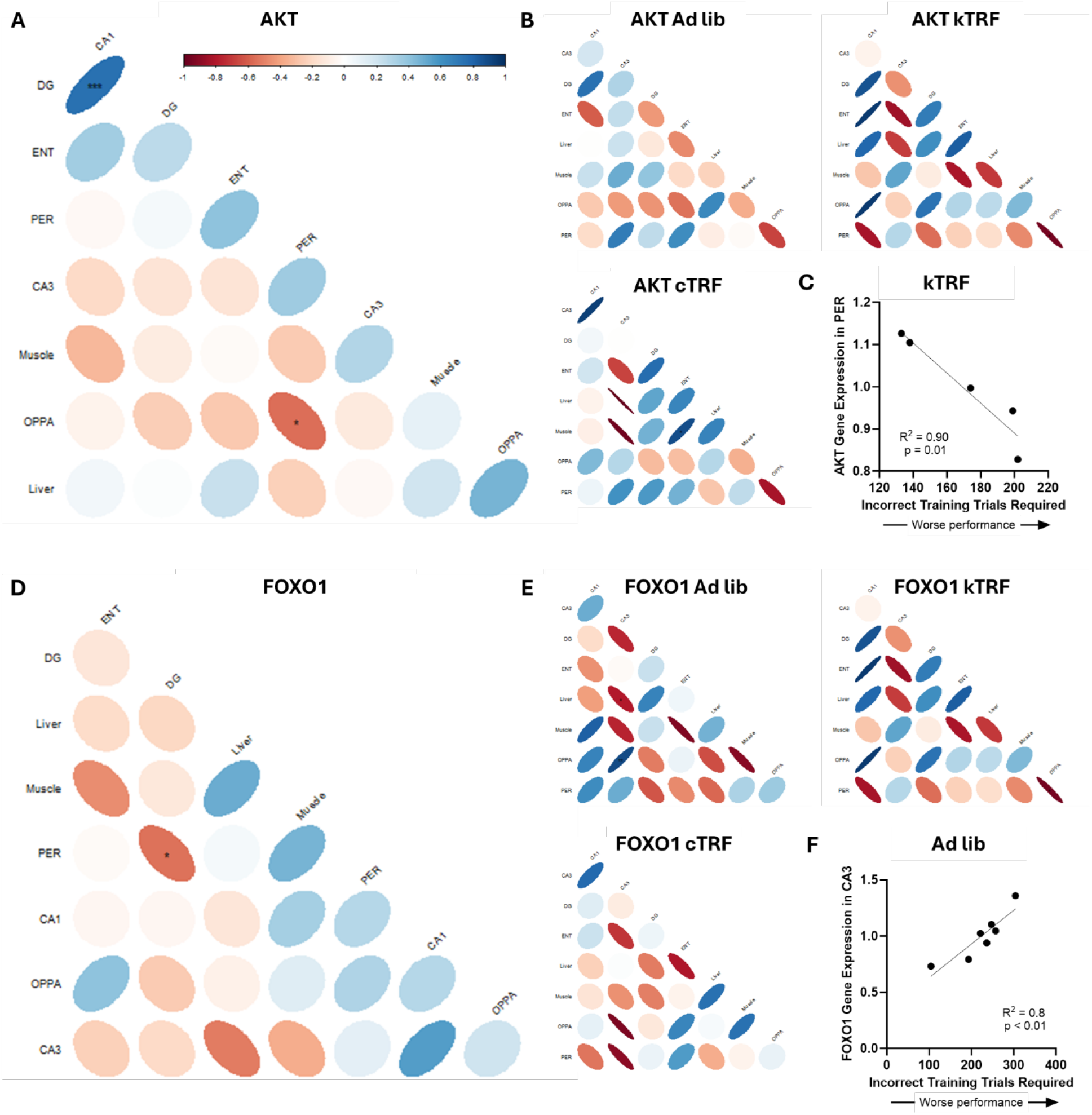
Cognitive performance on an object-place paired association (OPPA) task correlates with gene expression in a diet-specific manner. (A) Akt1 expression significantly correlated with behavioral performance when assessed across all rats within the perirhinal cortex (PER) only. (B) However, this correlation only remained significant for the kTRF diet group when analyzed separately, (C) with Akt1 expression significantly correlating with better behavioral performance in kTRF rats, though all groups demonstrated a trend towards a negative correlation. (D) Foxo1 expression did not correlate with behavioral performance when compared across all rats, though (E) again the effects were diet-specific, (F) and task performance significantly correlated with expression within CA3 of ad lib fed rats (and trended towards correlating with cTRF rats), with increased Foxo1 expression correlating with poorer cognitive performance.

Interestingly, the only significant correlation between gene expression levels across tissues was between CA1 and DG (r = 7.50; p < 0.001; all other comparisons p > 0.10). To investigate whether a certain diet group was driving these correlations, the analysis was rerun for each diet group independently (**Figure 2B**). The only group for which the OPPA performance and PER Akt1 expression remained significant was for kTRF rats (r = -0.095; p = 0.01; **Figure 2C**), though there was a trend for a correlation within cTRF (r = -0.82; p = 0.088) and ad lib (r = -0.67; p = 0.097). Within the cTRF rats, there was also a significant correlation between ENT and Muscle Akt1 expression (r = 0.89; p = 0.045), which trended for kTRF rats as well (r = -0.81; p = 0.09), but not ad lib fed rats (r = 0.19; p = 0.76).

Foxo1 expression across all rats, when all diet groups were combined, did not correlate with behavioral performance in any tissue (p > 0.12 for all comparisons; **Figure 2D**). However, expression of Foxo1 significantly correlated between DG and PER (r = -0.55; p = 0.02) across all rats, and there was a strong trend towards a correlation between CA1 and CA3 (r = 0.55; p = 0.06; **Figure 2E**). The relationship between Foxo1 gene expression and cognition was quantified within individual diet groups to see if dietary intervention affected these trends. Interestingly, there was a significant correlation between OPPA task performance and Foxo1 gene expression within the CA3 of ad lib fed rats (r = 0.90; p = 0.006; **Figure 2F**) as well as a trend in cTRF fed rats (r = -0.93; p = 0.069), but not in kTRF fed rats (r = -0.32; p = 0.61). In the ad lib fed rats, CA3 Foxo1 gene expression also significantly correlated with expression within the liver (r = -0.79; p = 0.03) and trended towards a correlation with DG (r = -0.75; p = 0.05). Together, these data suggest there is a relationship between Foxo1 and cognitive status under normal dietary macronutrient ratios in aged rats, but not while in nutritional ketosis.

Unlike Akt1 expression, expression of Foxo1 across tissues appeared to have a significant relationship across multiple tissue types, but only under ketogenic conditions. For kTRF rats, Foxo1 was significantly correlated between PER and ENT (r = 0.90; p = 0.04), CA3 and liver (r = -0.98; p = 0.02), CA1 and muscle (r = -0.998; p = 0.04), and ENT and muscle (r = -0.90; p = 0.04).

## Discussion

Previously, we demonstrated the positive impact of both a ketogenic diet (KD) and time restricted feeding (TRF) on cognitive performance and physical health [23,24], as well as neurotransmission related gene and protein expression within the brain [17,27]. The goal of the current study was to expand upon this work by investigating the effects of both KD and TRF on metabolism-related gene expression across tissues throughout the body. Herein we demonstrate tissue-specific effects of chronic TRF with or without a KD on PI3K/AKT signaling pathway gene expression in aged subjects. Not only are alterations in this pathway associated with aging and longevity, but also neurodegeneration and Alzheimer’s disease [28].

While the majority of genes within this pathway investigated were not significantly affected by the dietary intervention, expression of Akt1 itself was significantly altered in a tissue specific manner. Within the liver, KD significantly decreased expression. However, within the muscle, it appears that TRF had a greater effect on reducing expression. In addition to the gene expression investigations presented here, behavioral characterization of these subjects has been previously published [24], allowing for the investigation into the relationship between gene expression and cognitive performance. Regardless of diet group, there was a significant correlation between cognitive performance and Akt1 expression within the perirhinal cortex (PER). Despite a lack of differential gene expression across diet groups, the correlation with cognition remained significant only within the PER from kTRF rats when investigated by diet, with greater expression significantly correlating with better cognitive status. There was also a strong trend (adjusted p = 0.07) between cognitive performance and Akt1 expression within the liver.

While there have not been previous reports on changes in expression of the Akt1 gene across all these tissues following KD and/or TRF, especially in aged subjects, several others have looked at protein expression. Phosphorylated Akt (pAkt) protein level is decreased within both the hippocampus and liver of rats fed a ketogenic diet [29], though others have reported no change in pAkt within the liver [30]. A ketogenic-like diet enriched for ketogenic amino acids demonstrated a restoration of pAkt protein levels in muscle tissue following a high fat diet [31]. Our work, interpreted in the context of these protein data, indicates that the downregulation of Akt expression within both liver and muscle may be due to lack of need. In other words, sufficiently high AKT protein levels may have decreased the requirement for ongoing transcription in aged rats fed a KD.

However, no significant differences in Akt1 expression were observed in any of the brain tissues examined in the current study. It is possible that alterations in other neuronal pathways capable of modulating Akt1 expression compensated for systemic level changes in insulin signaling that modulated liver expression. For instance, neurotrophins such as brain-derived neurotrophic factor (BDNF) interact with the PI3K-AKT signaling pathway to moderate neuronal growth and plasticity [32]. Moreover, changes in Akt1 alone may not capture the full extent of AKT gene expression in the brain. Akt1, though important in peripheral insulin signaling, is expressed in the brain, but at weak levels; its expression increases dramatically in injured cells, which suggests that Akt1 may be more involved in cell survival and dealing with cellular damage in the brain [33]. Akt3, the dominant form of AKT in the brain, is associated with neurodevelopmental disorders and constitutes approximately half of total AKT protein in adult brains [34]. Each AKT isoform is associated with distinct functions, validating the need for further exploration into other AKT isoforms in the aged brain. Despite the lack of alterations in Akt1 expression within the brain, there was a significant correlation between PER Akt1 expression levels and the number of trials needed to acquire and object-in-place rule such that higher expression correlated with faster acquisition. It is possible that higher levels of Akt1 may be linked to more cell repair and synapse maintenance activity, which may lead to better maintenance and survival of the neurons in the PER. This correlation may have only been found in the PER due to its key roles in object recognition and as a pivotal gateway of communication between the HPC and the medial PFC [35].

While Akt expression did not change within the PER, Foxo1 expression in this brain region was significantly reduced in kTRF rats and trending towards a significant reduction in cTRF rats as well (adjusted p < 0.07), indicating these effects were likely an effect of TRF rather than KD. Foxo1 belongs to the forkhead box O-class (FoxO) subfamily of the forkhead transcription factors, a group suppressed via phosphorylation by the activation of the PI3k/Akt pathway. Foxo1 participates in a variety of signaling pathways and wide range of biochemical processes, including apoptosis, cell cycle transition, DNA repair, muscle growth and atrophy and much more [36]. Thus, decreased expression of this gene within the brain may indicate several things. Firstly, Foxo1 plays a role in promoting gluconeogenesis [37] and influences how neurons manage glucose [38]. Secondly, Foxo1 impacts lipid synthesis and storage [39]. Thus, transitioning from a glucose based to a ketone body based metabolic state within the brain could reasonably result in decreased Foxo1 requirement to maintain these now underutilized processes. Foxo1 is also critical for antioxidant defense. Thus, decreased gene expression may be indicative of less oxidative stress requiring protection. Additionally, Phosphorylated Foxo1 protein is targeted for proteasome degradation [40]. Thus, it is possible that decreased Foxo1 gene expression may be due to decreased need for additional Foxo1 protein due to lower turnover linked to less frequent AKT signaling. This suggests that Foxo1 protein may remain unphosphorylated and active more frequently in TRF groups, which may suggest less activity of AKT linked to insulin signaling.

Interestingly, there was only a significant relationship between Foxo1 gene expression and cognitive status in control-fed rats, but not ketogenic-fed rats. Within CA3, higher levels of Foxo1 gene expression were correlated with worse ability to acquire a biconditional association rule in ad lib-fed rats. Moreover, higher levels of Foxo1 gene expression within CA3 also significantly correlated with lower levels of expression with the liver in ad lib-fed rats. These effects were not observed in KD fed rats. This could indicate that Foxo1 plays a role in cognitive function under normal conditions, but not under nutritional ketosis. Alternatively, this could indicate that age-related disruptions in Foxo1 signaling are ameliorated in the aged subjects fed a KD.

The data presented herein differ from previous reports on the effects of KDs on PI3K/AKT expression in several ways. Firstly, these rats have been keto-adapted for a significant portion of their entire adult lives, which is rarely the case in existing literature. Secondly, this version of a ketogenic diet includes MCT oil, which is known to be significantly more beneficial in many ways relative to other long chain fat based diets (especially those using lard as the primary fat source added to standard chow) [41–43]. These specific methods of dietary implementation may explain why we observed significantly decreased AKT gene expression, whilst others have shown increased protein expression. There are several key limitations to this work as well. Firstly, caution should be exercised as to not over interpret gene expression data, wherein decreased expression can indicate both an impaired ability to generate required proteins or a decreased need for the continued transcription of a gene. Secondly, this work only included male subjects. The inclusions of females, who have notable metabolic differences across the lifespan [44], may result in differential gene expression. However, this work incorporated near life-long dietary manipulation and investigation of tissues from aged subjects, and thus still provides valuable insight into dietary practices implemented in advanced age.

## Acknowledgements

We thank Lijiang Guo and Stephanie Dickinson of the UAB Nathan Shock Center Data Analytics Core and Indiana University School of Public Health-Bloomington Biostatistics Consulting Center for support with data analysis.

## Statements and Declarations

The authors declare that they have no conflicts of interest.

